# DeepSEA: an alignment-free explainable approach to annotate antimicrobial resistance proteins

**DOI:** 10.1101/2024.06.11.598242

**Authors:** Tiago Cabral Borelli, Alexandre Rossi Paschoal, Ricardo Roberto da Silva

**Affiliations:** Computational Chemical Biology Laboratory, Department of BioMolecular Sciences, School of Pharmaceutical Sciences of Ribeirão Preto, University of São Paulo, Ribeirão Preto 14040-900, Brazil; NPPNS, Department of BioMolecular Sciences, School of Pharmaceutical Sciences of Ribeirão Preto, University of São Paulo, Ribeirão Preto, 14040-900, Brazil; Cellular and Molecular Biology Program, Department of Cellular and Molecular Biology of Ribeirão Preto, School of Medicine, University of São Paulo, Ribeirão Preto, 14049-900, Brazil; Bioinformatics and Pattern Recognition Group (Bioinfo-CP), Department of Computer Science (DACOM), The Federal University of Technology – Paraná (UTFPR), Cornélio Procópio, Brazil; Rosalind Franklin Institute, Harwell Science and Innovation Campus, Didcot OX11 0QS, UK

## Abstract

Antimicrobial resistance (AMR) is one of the most concerning modern threats as it places a greater burden on health systems than HIV and malaria combined. Current surveillance strategies for tracking antimicrobial resistance (AMR) rely on genomic comparisons and depend on sequence alignment with strict similarity cutoffs of greater than 95%. Therefore, these methods have high false-negative error rates due to a lack of reference sequences with a representative coverage of AMR protein diversity. Deep learning (DL) has been used as an alternative to sequence alignment, as artificial neural networks (ANNs) can extract abstract features from data and therefore limit the need for sequence comparisons. Here, a convolutional neural network (CNN) was trained to differentiate antimicrobial resistance proteins from nonresistance proteins and functionally annotate them in nine resistance classes. Our model demonstrated higher recall values (>0.9) than the alignment-based approach for all protein classes tested. Additionally, our CNN architecture allowed us to investigate internal states and explain the model classification regarding protein domain feature importance related to antimicrobial molecule inactivation. Finally, we built an open-source bioinformatic tool that can be used to annotate antimicrobial resistance proteins and provide information on protein domains without sequence alignment.

## Background

Penicillin discovery (Fleming, 1929) enabled the treatment of harsh bacterial infections such as syphilis. This approach ensures the safety of patients through invasive or immunosuppressive procedures, which promotes a substantial increase in human life expectancy (Yoshikawa, 2002). However, either producing or resisting antimicrobials are competitive mechanisms in microorganisms’ evolution (Skinnider et al., 2020), and the overuse and misuse of these drugs have been selecting bacterial variants with genotypes that confer resistance. This increasing-dosage increasing-resistance dynamics initiated the *Antibiotic Crisis* (Neu, 1992; Rossolini et al., 2014; Wright, 2015), a *Red Queen* race (Leigh Van Valen, 1973) - a term borrowed from Alice Through the Looking-Glass to represent positive feedback loops - that resulted in 1.27 million deaths directly caused by antimicrobial resistance (AMR) in 2019 (Murray et al., 2022).

Given the aforementioned, there is a need to develop new drugs to fight resistant bacteria and, most importantly, surveillance strategies to track AMR (Djordjevic et al., 2023). The current approach is based on sequence alignment to identify resistance genes and proteins. In the alignment-based model, candidate and reference sequences are considered homologous if the sequence similarity exceeds a given threshold. Dedicated databases and tools based on antimicrobial susceptibility tests (ASTs) (Hasman et al., 2014) have different cutoffs to annotate query sequences as resistant proteins. For instance, Resistance Gene Identifier (RGI) (Alcock et al., 2020) uses curated bitscore values of each reference protein stored in its database, whereas AMRFinderPlus (Feldgarden et al., 2019) and ResFinder (Florensa et al., 2022) use raw similarity and coverage values.

The dependency on reference sequences for alignment-based models constrains surveillance to rediscover the same proteins/genes and to have a high number of false negatives (resistant proteins classified as nonresistant) (Arango-Argoty et al., 2018), as most tools use strict cutoffs above 80% (Florensa et al., 2022) or 95% (Pal et al., 2016) of similarity to annotate a protein as resistant. Deep learning-based models have emerged as a solution to this issue, as neural networks can learn complex and nonlinear rules from data and use them to annotate proteins without direct sequence alignment to known references (Bileschi et al., 2022). In this regard, the pioneers DeepARG (Arango-Argoty et al., 2018) and HMD-ARG (Li et al., 2021) achieved remarkable results in multiclass task classification of resistance proteins with lower false negative rates than alignment-based alternatives. However, these tools either need protein alignment in some stages of their workflow or need a complex approach to make the output explainable.

In this work, we present a study on the ability of the convolutional neural network (CNN) to annotate resistance proteins. Our CNN model had an equivalent performance to the cutting-edge protein language model from the Evolutionary Scale® (https://www.evolutionaryscale.ai/) and outperformed the alignment-based approach. The CNN was able to classify proteins across nine classes and differentiate them from nonresistance proteins with recall (true positives / relevant elements) values above 0.95. We also presented an algorithm to process CNN’s neurons firing partners that explain the CNN decision-making. Finally, we wrap the model and the algorithm in a command-line interface tool to provide insightful outputs for users with a biological background.

## Methods

### Database construction

For model training, hyperparameter optimization, and evaluation, we selected the Non-redundant Comprehensive Database (NCRD) (Mao et al., 2023a). The NCRD was built to provide more reference sequences for alignment-based annotation of antimicrobial proteins. The NCRD developers collected consolidated resistance proteins from the Antibiotic Resistance Genes Database (ARDB) (Liu and Pop, 2009), Comprehensive Antibiotic Resistance Database (CARD) (Alcock et al., 2020), and Structured Antibiotic Resistance Genes (SARG) (Yin et al., 2018) and expanded the database using DIAMOND (with the parameters E-value ≤ 1×10-5, Query Coverage HSP ≥ 90%, and Percentage of Positive Positions ≥ 90%) to find homologous proteins from Non-redundant Protein Database (https://www.ncbi.nlm.nih.gov/refseq/about/nonredundantproteins/) and Protein Bank Database (https://www.rcsb.org/). For this work, we selected the NCRD95, a database version in which the sequence similarity was limited to 95% by CD-HIT (Li and Godzik, 2006).

The NCRD95 was converted from FASTA to table format via SeqKit 2.8.1 (Shen et al., 2024), and the FASTA identifiers were inspected to retrieve information on the content of antimicrobial resistance protein classes. Cumulative class curves were employed to (i) determine the optimal number of protein resistance classes that minimize data imbalance while maximizing the database size and (ii) eliminate less representative protein subclasses within each class.

To build the non-resistance class (NonR), we downloaded reviewed proteins (SwissProt proteins) that are not associated with antimicrobial resistance (Arango-Argoty et al., 2018; Li et al., 2021; Wu et al., 2023) with the command *(taxonomy_id:2) NOT (keyword: KW-0046)* on the Uniprot website and applied CD-HIT to limit protein similarity to 95%. Finally, DIAMOND (Buchfink et al., 2015) (with coverage > 80%, E-value ≤ 1×10-3, and max-target-seqs = 1) was used to align SwissProt against NCRD95 proteins. The number of proteins that significantly aligned despite not being associated with antimicrobial resistance, according to SwissProt (approximately 4600), was kept and used to randomly sample the proteins that were not aligned to NCRD95. Therefore, the NonR class is composed of proteins with no relevant sequence identity to antimicrobial resistance proteins in our dataset.

### Model architecture, training, and evaluation

We used the *train_test_split* function from scikit-learn 1.4.2 to split the final dataset into training and holdout test sets at an 80/20 ratio, stratified according to the original protein class distribution. The training set was used for hyperparameter (Table 1) optimization with the Keras Hyperband (https://keras.io/keras_tuner/api/tuners/hyperband/) heuristic algorithm and 5-fold cross-validation (five random stratified split points). The hold-out test was used for model evaluation and benchmarking. For reproducibility, the *random_state* (seed) parameter was set to 42 for all split steps performed in this work.

**Table 1.**
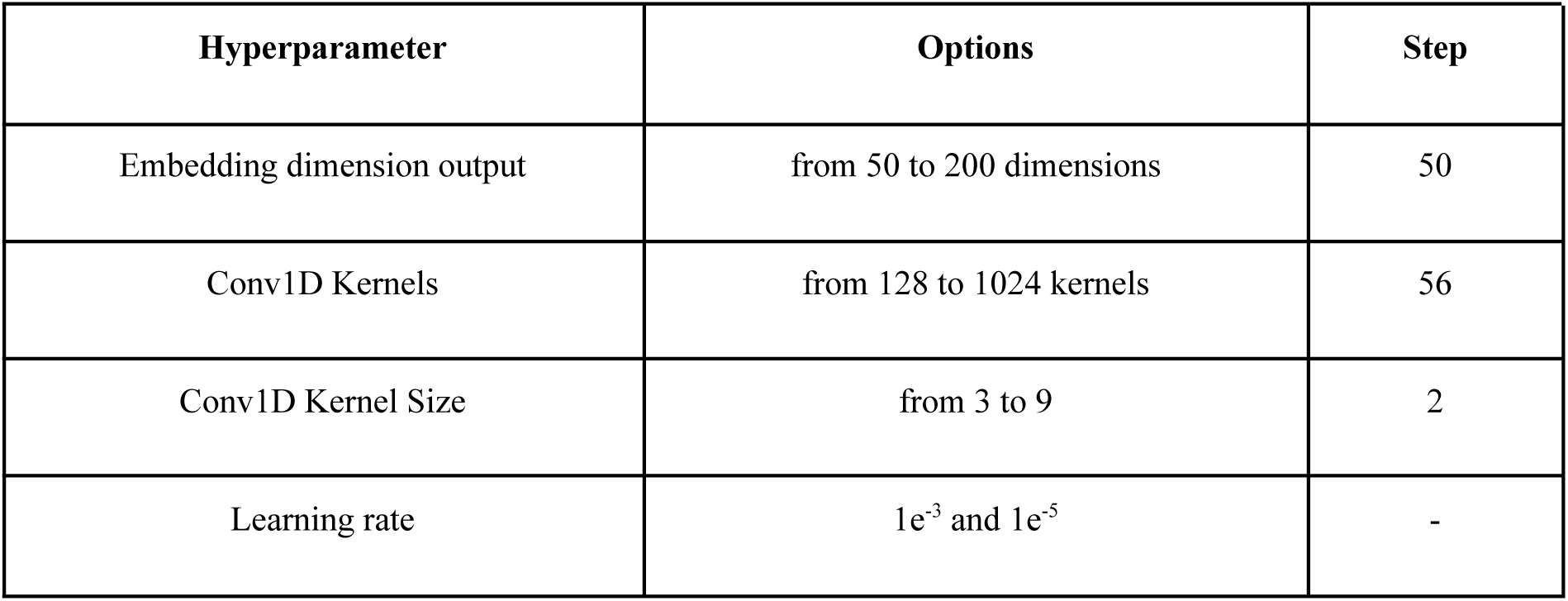
Hyperparameter space.

To establish a baseline for benchmarking, we used Transformers 4.49 (Wolf et al., 2020) and PyTorch 2.6 (Paszke et al., 2019) to fine-tune an Evolutionary Scale Model 2 (ESM2) (Lin et al., 2023) from a protein language model of general-purpose to an antimicrobial resistance protein classifier. Additionally, RGI and AMRFinderPlus outputs were manually inspected and mapped from antimicrobial molecule names to broader classes, and results below their respective thresholds were considered nonresistant proteins.

We used TensorFlow 2.15 and Keras 2.15 to design an end-to-end convolutional neural network model that received input batches of raw amino acid sequences instead of multiple sequence alignment files. Our architecture contains 3 blocks: (i) a data processing block that contains a text2vec as the input layer and a subsequent embedding layer, (ii) a feature extraction block: composed of four 1D convolutional layers interpolated with dropout layers used for regularization, and (iii) a classification block with global average pooling and dense layers.

Reference proteins from the same classes were retrieved from the reference catalog of the National Database of Antibiotic-Resistant Organisms (NDARO) and used to evaluate our model on an independent dataset. The DIAMOND (E-value ≤ 1×10-3, and max-target-seqs = 1 and coverage == 100%) was used to align them against your training set to reveal the distribution.

### Feature extraction

*Algorithm 1* describes how we access the matrices within our CNN and process them (Additional File 1; Supplementary Figure 1). First of all, a protein sequence is converted to tensor format to fit input requirements (line 2), and then the CNN model is applied (line 4) to create a probability distribution from which the most probable class index is retrieved (line 6). The dense layer contains weight vectors to calculate logits before probability distribution, and the TensorFlow/Keras frameworks allow access to these vectors. Therefore, in lines 8 and 9, the algorithm retrieves the weights vector for the class predicted by the model. In line 11, a submodel is temporarily created by removing the global average pooling and dense layers from our CNN. This new model’s output is a matrix with features extracted by convolution. The protein sequence is used by the submodel to obtain and retrieve the feature matrix (lines 13 and 14). Finally, the feature matrix and the weights vector are multiplied (line 16) to highlight the most important features in a normalized (line 18) vector with the same length as the imputed protein.

To check if the final weight vector matches the biological properties of the resistance proteins, we combined Algorithm 1 and multiple sequence alignments of enzymes from your holdout test set. For visual clarity, we first clustered your enzymes by sequence similarity (up to 90%) and minimal coverage equal to 90%, and the largest cluster of each protein class was selected to be aligned via MAFFT 7.5 (Kuraku et al., 2013). To highlight protein regions of greater weights, we represent the multiple sequence alignments as matrices where the proteins are replaced by their corresponding vectors and plotted as heatmaps. The gaps were filled with zeros. Finally, InterproScan 5.0 (Blum et al., 2021; Jones et al., 2014) was applied to annotate important protein motifs, and Shannon entropy was calculated for each position to reveal conserved regions. For motif analyses, we chose the proteins used as a cluster reference by CD-HIT.

#### Algorithm 1

**Extract weights**

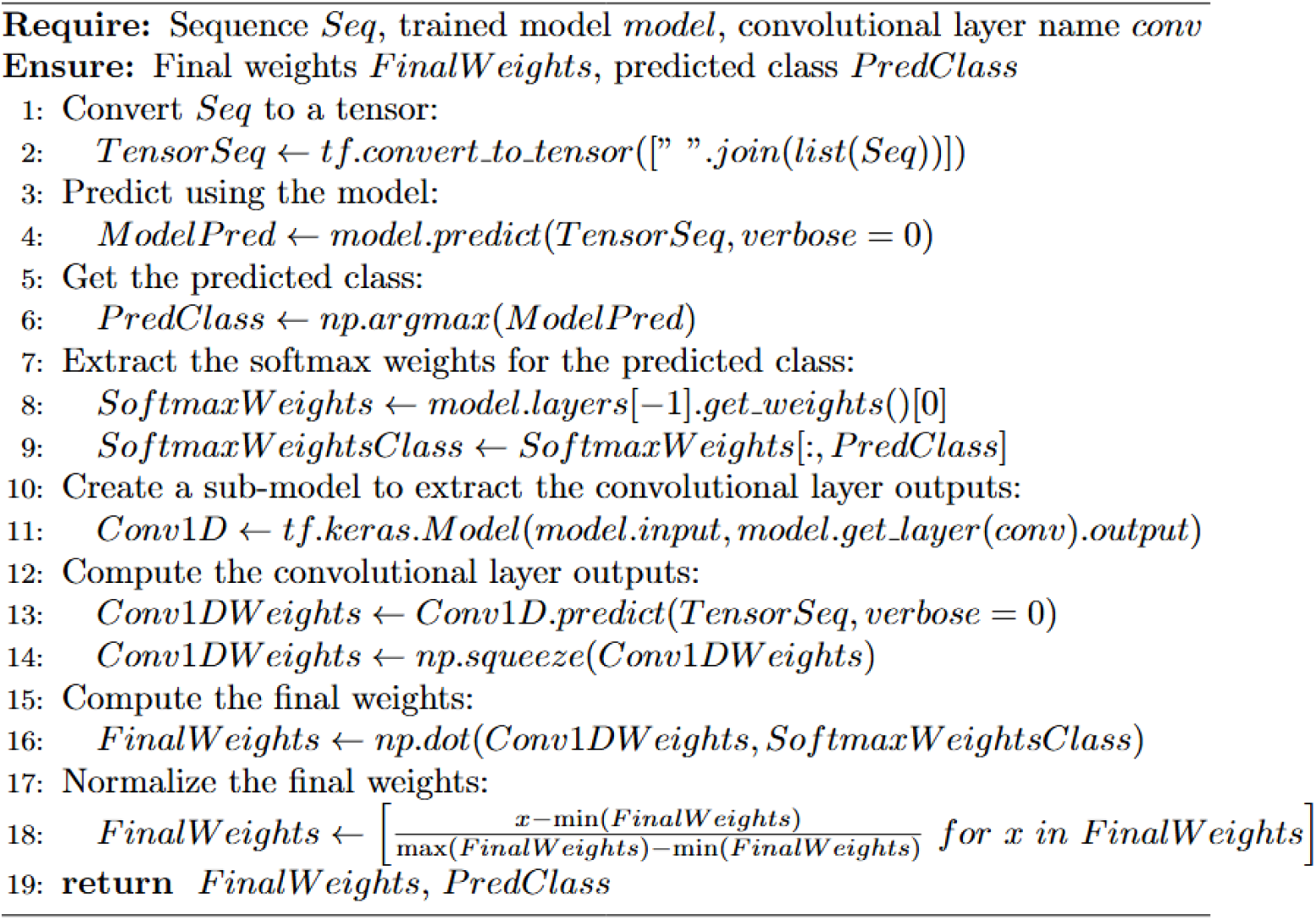

### CNN cluster ability

To assess cluster ability, we leveraged the information embedded in the global average pooling layer. As this layer works as a bridge between feature extraction and the final classification made by the dense layer, its internal state represents each protein imputed as a multidimensional vector with summarized features extracted upstream. Therefore, we designed an algorithm (*Algorithm 2*) that removes the dense layer and makes the model output the global average pooling vectors.

#### Algorithm 2

**Dimension reduction**

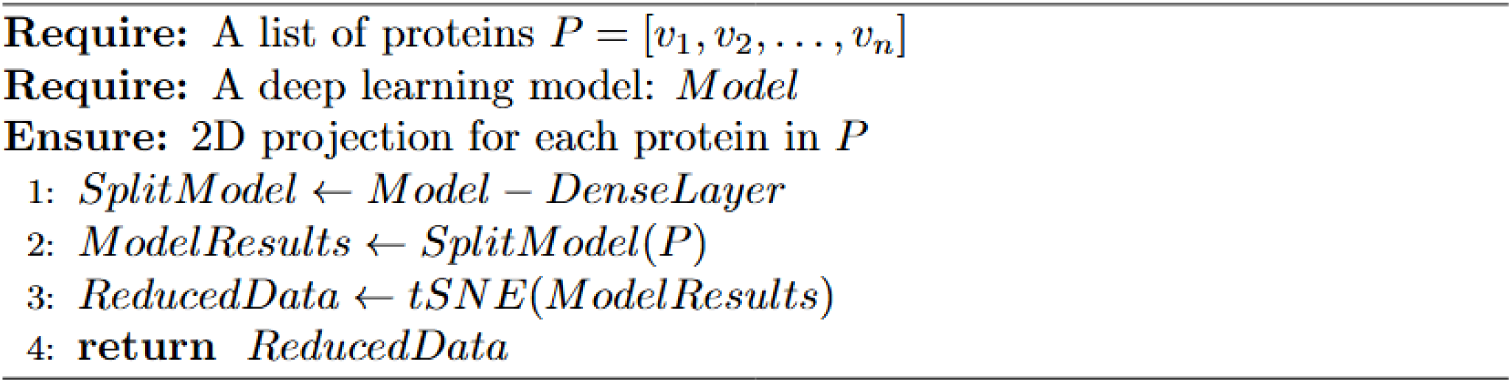

*Algorithm 2* was applied on the holdout test set since the model has no knowledge of these proteins and the resulting vectors (Additional File 1; Supplementary Figure 1). The resulting vectors were then concatenated into a matrix and subsequently reduced to two dimensions through t-distributed stochastic neighbor (t-SNE) from scikit-learn 1.4.2 with the following parameters: t-SNE learning rate set to *auto*, iterations = 1000, and perplexity = 50. The original index order from the holdout test was kept. Thus, the reduced matrix contained the respective classes of the protein.

### Evaluation metrics

To address biases stemming from uneven data, we incorporated class weights to avoid errors in model performance assessment linked to an imbalance in protein class sizes. The calculation of precision (Equation 1), recall (Equation 2), the f-score (Equation 3), and the categorical cross-entropy error (Equation 4) incorporates the weight of each protein class during training.

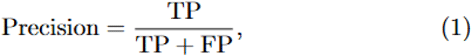

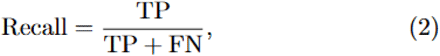

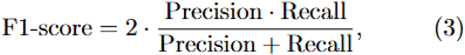

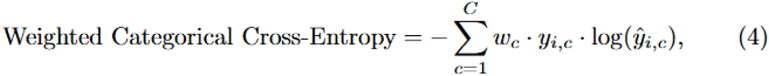

where:

- *TP* is the number of true positives,
- *FP* is the number of false positives,
- *TN* is the number of true negatives,
- *FN* if the number of false negatives,
- *C* is the number of classes,
- *w_c_* is the weight for class *c,*
- *y_ic_* is the indicator if class c is the correct label for sample *i*,
- *y_ic_* is the predicted probability by the neural network that sample *i* belongs to class *c*.

### Class weights calculation

The class weights (Additional File 2; Supplementary Table 1) were computed using scikit-learn’s compute_class_weight 1.4.2 (Equation 5), which assigns weights inversely proportional to class frequencies in the dataset to address imbalance.

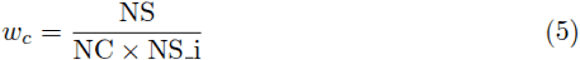

Where:

- NS: Total number of samples in the dataset
- NC: Number of unique classes
- NS_i: Number of samples in class *i*

## Results

### Dataset curation and the lack of reference sequences

CARD and National Database of Antibiotic-Resistant Organisms (NDARO) are the two largest public databases used as sources of reference sequences for functional annotation of antimicrobial resistance proteins. Since the CARD requires experimental validation, it has been widely used in several studies. (Arango-Argoty et al., 2018; Li et al., 2021; Mao et al., 2023b; Z. Wang et al., 2021; Wu et al., 2023). Even part of the NDARO database is composed of CARD. However, since a protein can be annotated only if it is highly similar to a reference in the CARD, researchers are constantly rediscovering variations of the same resistance proteins.

We investigated the CARD database updates (Additional File 1; Supplementary Figure 2A) to check the rate at which novel reference proteins were added over the last few years. Our findings revealed that, as expected, the total number of proteins reached a plateau, while antimicrobial resistance has been increasing (Murray et al., 2022) and making infection treatment ineffective (Liu et al., 2016). The NCRD95 database extended CARD content using homology search and was, therefore, selected for this work.

Originally, NCRD95 contained 29 antimicrobial resistance protein classes (Additional File 1; Supplementary Figure 1B) from 10603 (multidrug efflux pumps) to 2 (diaminopyrimidine) examples. This high degree of imbalance compromises training, as classes with few examples may not provide enough information for generalization (B. Wang et al., 2021). Therefore, we plotted a cumulative curve (Additional File 1; Supplementary Figure 1C) and selected the 10 most abundant classes (Additional File 1; Supplementary Figure 1D), as beyond this point, more classes did not substantially increase the total number of proteins. For instance, increasing protein classes from 10 to 15 would add only 1103 proteins (a 3% increase) to our final database. We also performed the same process for each class individually to eliminate less representative subclasses and ensure internal homogeneity.

Data leakage among classes (i.e., classes that share information) that involve multidrug resistance proteins has also been addressed. This class consists mainly of efflux pumps that extrude different antimicrobial molecules from bacterial cells, which means that the same protein may be classified into two classes depending on the experimental design. For example, we found RND efflux pumps in the multidrug, beta-lactam, and macrolide classes. Therefore, we removed the multidrug class and all efflux pumps found in our dataset, regardless of the protein class. Finally, we constructed a homology-based dataset composed of nine antimicrobial resistance proteins plus one class of nonresistance proteins (Additional File 1; Supplementary Figure 1D).

### Deep learning model evaluation

The Keras Hyperband algorithm took 83 rounds of heuristic search to optimize the set of hyperparameters, which yielded a CNN model with the lowest loss value possible (Additional file 2; Supplementary Table 2). However, low loss values must be taken cautiously in deep learning research since neural networks can learn the intrinsic noise of the training set rather than the meaningful signal. In the protein research field, this may occur when there are copies of proteins shared by training and test sets, so the model would recognize idiosyncratic features instead of general features. We mitigated the risk of overfitting resulting from data leakage by restricting the sequence similarity between the training and test sets to a maximum of 95%

Another precaution taken was the split point that created the training set. Although randomized, the process may lead to a biased model if the proportion of classes between training and test is not respected. We addressed this issue by cross-validating our model with 5 stratified sub-splits of our training set. The convergence curves of cross-validation (Additional File 1; Supplementary Figure 3A), final model training (Additional File 1; Supplementary Figure 3C and 3D), and evaluation metrics (Table 2) on the holdout test set showed no clear trends of overfitting regardless of the training subset.

**Table 2.** Cross-Validation Results.

| Class | Mean precision | Mean recall | Mean F1-score |
| --- | --- | --- | --- |
| MLS | $0.96 \pm 0.05$ | $0.95 \pm 0.02$ | $0.95 \pm 0.02$ |
| NonR | $0.83 \pm 0.02$ | $0.97 \pm 0.02$ | $0.89 \pm 0.01$ |
| Aminoglycoside | $0.96 \pm 0.03$ | $0.95 \pm 0.01$ | $0.96 \pm 0.02$ |
| Beta-lactam | $0.99 \pm 0.01$ | $0.96 \pm 0.01$ | $0.97 \pm 0.01$ |
| Chloramphenicol | $0.99 \pm 0.01$ | $0.98 \pm 0.01$ | $0.99 \pm 0.01$ |
| Glycopeptide | $0.99 \pm 0.01$ | $0.99 \pm 0.04$ | $0.99 \pm 0.002$ |
| Macrolide | $0.98 \pm 0.01$ | $0.87 \pm 0.07$ | $0.92 \pm 0.04$ |
| Phosphonic Acid | $1.0 \pm 0$ | $0.99 \pm 0.03$ | $0.99 \pm 0.015$ |
| Rifamycin | $0.99 \pm 0.01$ | $0.98 \pm 0.01$ | $0.99 \pm 0.003$ |
| Tetracycline | $0.98 \pm 0.03$ | $0.98 \pm 0.01$ | $0.98 \pm 0.01$ |

We use the holdout test set to compare deep learning and protein alignment on antimicrobial resistance protein annotation. To represent the alignment-based approach, the state-of-the-art tools Resistance Gene Identifier (RGI) 6.0.3 (Alcock et al., 2020) and AMRFinderPlus 4.0.19 (Feldgarden et al., 2019) were selected. On the deep learning approach, our CNN and the fine-tuned ESM2. To equalize the number of classes, proteins not classified as resistant by RGI or AMRFinderPlus were considered nonresistant.

The deep-learning-based approach (Figures 1A and 1B) presented high values of recall (>0.95) across antimicrobial resistance protein classes, although their performance slightly decreased in the precision of the nonresistance classes. On the other hand, alignment-based tools (Figure 1C and Figure 1D) only achieved comparable values of precision and recall in cases where there are sequences that serve as references for annotating proteins.

**Figure 1.**
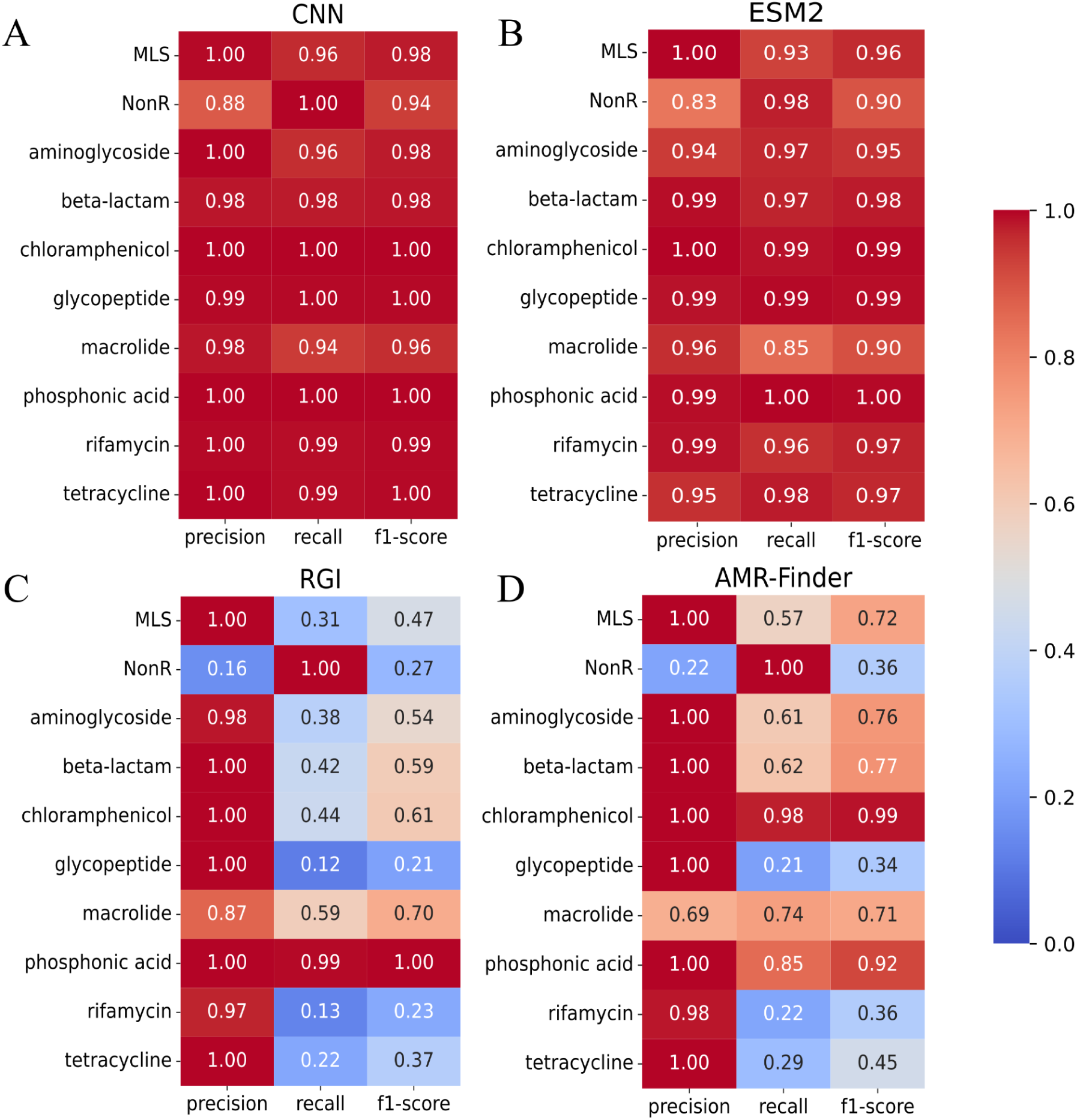
Classification reports. Benchmarking of CNN (A), ESM2 (B), RGI (C), and AMR-Finder (D) on the holdout test set. The metrics were calculated with class weight values to avoid bias due to class imbalance.

Confusion matrices showed in detail the misclassifications of each tool in our benchmarking. CNN and ESM2 (Figure 2A and Figure 2B) presented a few antimicrobial resistance proteins annotated as nonresistance proteins. RGI and AMR-FinderPlus (Figure 2C and Figure 2D), on the other hand, presented more false negative results, mostly in the glycopeptide class.

**Figure 2.**
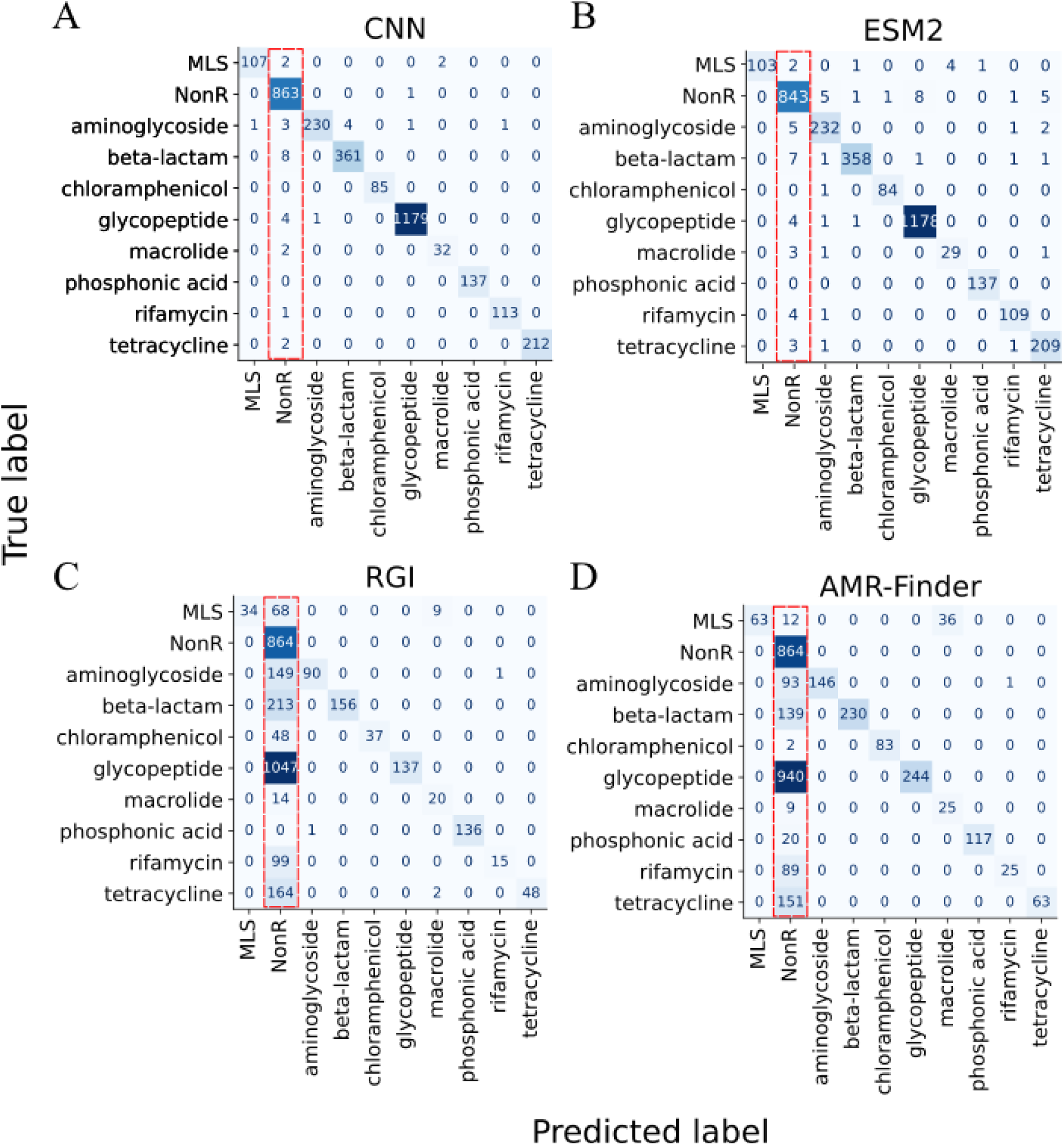
Confusion matrices of CNN (A), ESM2 (B), RGI (C), and AMR-Finder (D) on the holdout test set. The red-dashed squares highlight the false negative results of each benchmarked tool.

### Generalization in out-of-distribution data

Antimicrobial resistance proteins are limited in number due to the sequence alignment drawbacks mentioned earlier. For instance, CTX-M beta-lactamases share 94% of amino acid identity (ur Rahman et al., 2018). Such a lack of diversity may impose biases into deep learning models, which would perform well only on proteins identical to their training set but poorly on dissimilar proteins (Bernett et al., 2024). Therefore, we investigated the limits of generalization of our CNN regarding the sequence similarity using our holdout test set and the NDARO.

DIAMOND alignments showed that approximately 68% (2292 out of 3352 proteins) of the holdout test set aligned to the training set with an identity range of 29% to 95%. This indicates that our CNN is not limited to identical variants of the training set; otherwise, the model would demonstrate a significant decrease in its performance. CNN’s performance on the remaining 32% (1060 proteins) corroborates this hypothesis, as the values of precision, recall, and f1-score remained high (Additional File 1; Supplementary Figure 4A). In fact, for chloramphenicol and phosphonic acid, they remained equal to one. The decrease observed in the precision of non-resistant proteins resulted from the few false negatives from the other classes.

Our CNN yielded similar results on NDARO proteins. From the 5959 resistant proteins, 5726 aligned to the training set, and 5654 were correctly classified (Additional File 1; Supplementary Figure 4B). From the set of proteins that did not align to our training set, DeepSEA misclassified 42 proteins from beta-lactam (Additional File 1; Supplementary Figure 4C). A manual inspection revealed that the majorly of these 42 proteins are subclasses absent in our training set, such as VEB (n=11), CFX (n=12), and MUN (n=6) beta-lactamases (Additional File 1; Supplementary Figure 4D). Indeed, only the subclass OXA (n=1) contains examples in our training and was wrongly assigned as nonresistant. Together, these results provided evidence of true generalization as our CNN annotated out-of-identity-distribution proteins, as long as there are examples in the training set.

### Model explainability

The results exposed above led us to investigate the latent space produced by the feature extraction block to reveal which protein features the CNN learned to yield such high performance. The global average pooling layer summarizes the features extracted by the convolutional layers. Therefore, we applied *Algorithm 2* to make the model return these vectors instead of classifying proteins and reducing them with t-SNE. Our CNN separated the holdout test set according to protein classes (Figure 3) with a few misclassifications, indicating that the model successfully differentiated sets of weights for each protein class, which shows the model’s ability to distinguish resistant from nonresistant proteins and predict the correct resistance classes.

**Figure 3.**
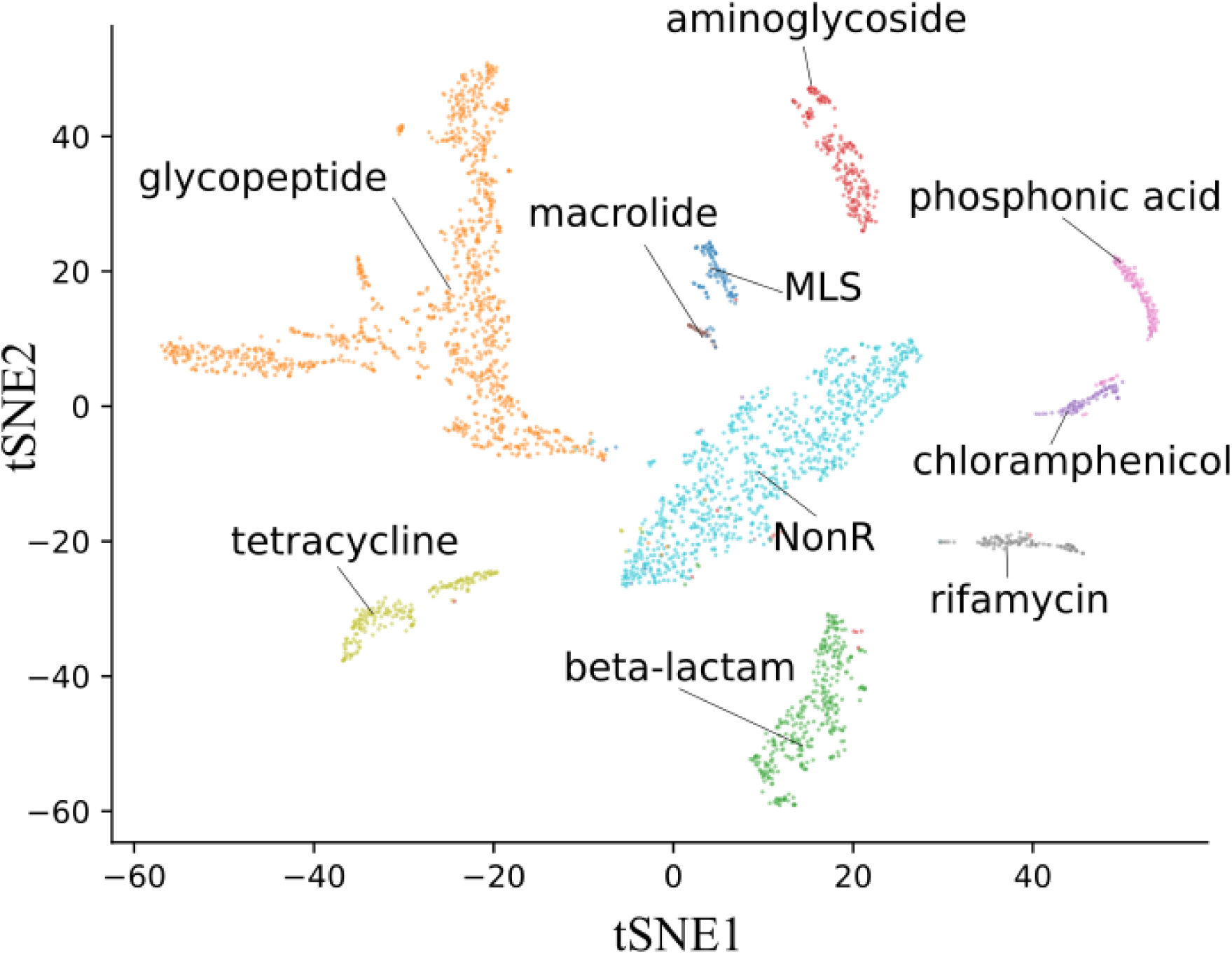
CNN class clusters. The t-SNE plot is made of the reduced weight matrix retrieved from the global average pooling layer as described in Algorithm 2 (see Methods).

As these results indicate that there are diagnostic features learnt from each protein class, we searched for them in the feature matrices produced by the convolutional layers, and investigated if they could be associated with biological functions. Briefly, *Algorithm 1* highlights important features by dot-producting the weights from the dense layer and the feature matrix for a given imputed protein. Pairing the resulting vector with the original protein sequence reveals residues of importance. Also, we overlapped the feature vectors and protein domains annotated by InterproScan 5 to add biological meaning.

Our results showed that CNN selected conserved regions (determined from multiple sequence alignments) as information hotspots for classification. For instance, class-A beta-lactamases (Figure 4A) alignment presented three main regions where CNN neurons fired to predict that class of enzymes, all of which are within the large beta-lactamase signature (IPR000871). The first hotspot contains the class-A active site (IPR023650) where the catalytic residues (S--K) are located (Ghuysen, 1991; Joris et al., 1988a). The last hotspot includes the binding site GTK, another conserved and diagnostic subdomain of class-A beta-lactamases.

**Figure 4.**
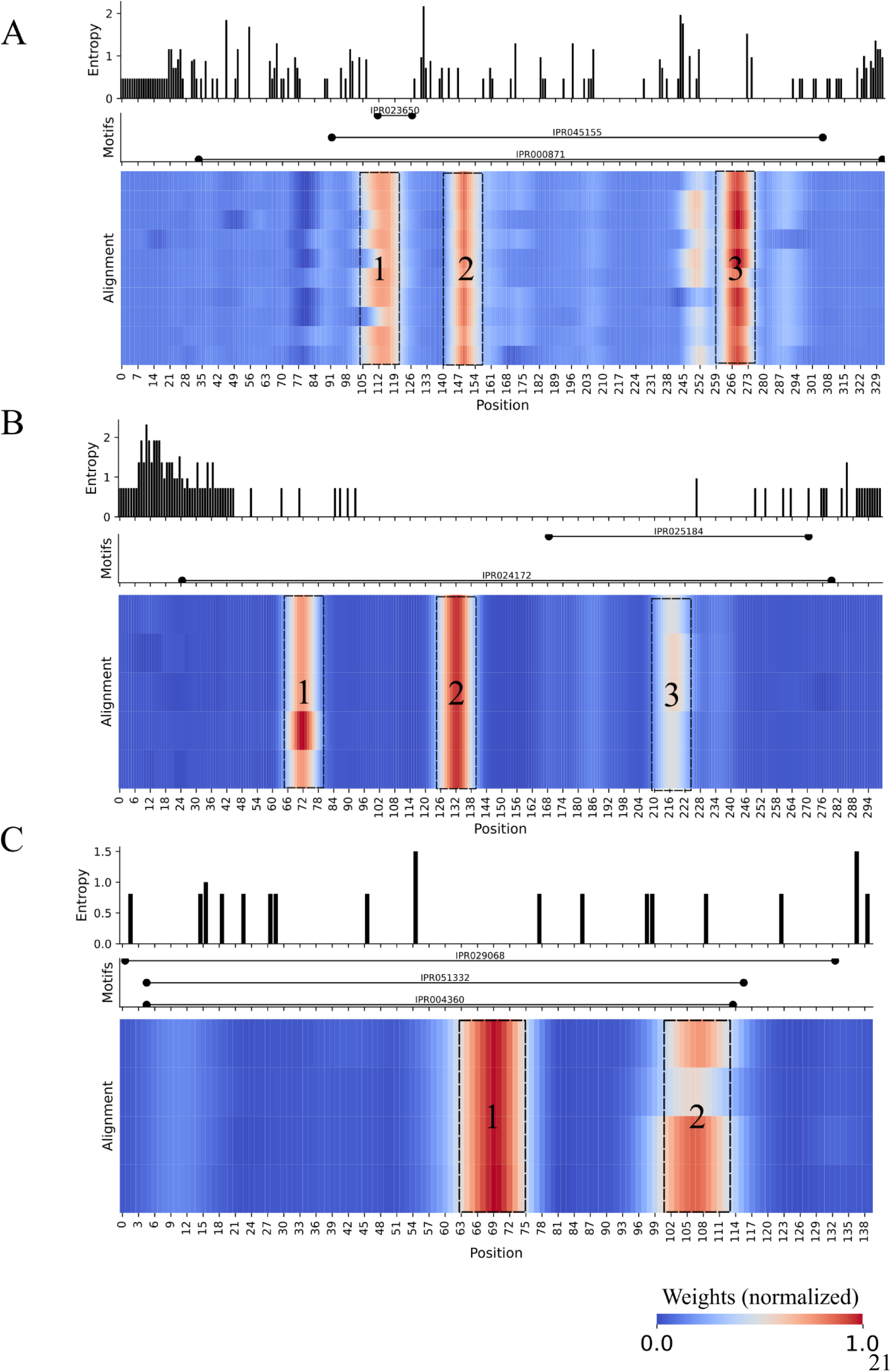
CNN neurons firing patterns for resistance proteins from the beta-lactam (A), aminoglycoside (B), and phosphonic acid (C) classes. Each plot is divided into a bar plot with Shannon entropy values, a line plot with protein domain extensions found by Interproscan 5, and the CNN weights spread over a multiple sequence alignment of a cluster of proteins selected by CD-HIT (see Methods). The black-dashed squares highlight the most important regions (hotspots) for classification.

The aminoglycoside-modifying enzyme AadA alignment (Figure 4B) also has three important conserved regions that belong to the adenylyltransferase AadA/Aad9 domain (IPR024172). The last hotspot is in the adenylyltransferase AadA, C-terminal domain (IPR025184), and the first belongs to the nucleotidyltransferase domain from the superfamily of proteins in which adenylyltransferases are included. Fosfomycin thiol transferase (FosA), the example of the phosphonic acid (or fosfomycin) class (Figure 4C), has two hotspots in conserved domains related to fosfomycin resistance: fosfomycin resistance protein (IPR051332), glyoxalase/fosfomycin resistance/dioxygenase domain (IPR004360) and glyoxalase/bleomycin resistance protein/dihydroxybiphenyl dioxygenase (IPR029068).

Given these results, we enquired whether our model captured information from protein regions related to antimicrobial molecule inactivation in the cases of aminoglycoside and fosfomycin-resistant proteins. To investigate this, we selected enzymes in complex with antimicrobials from the Protein Data Bank PDB (Table 3), applied the same feature extraction method, and compared our results with PDB functional annotations. CTX-M15 beta-lactamase (PDB: 7TI0) (Figure 5A) hotspots contain residues annotated as binding sites. Specifically, the residues S--K in which the serine (S45) residue nucleophilically attacks the lactam ring and initiates the structural modification that leads to beta-lactam molecule inactivation (Biswal et al., 2023; Ghuysen, 1991; Judge et al., 2023; Majiduddin et al., 2002). Additionally, the last hotspot contains the residues K209, T210, and G211, which bind and place the beta-lactam molecule in the catalytic pocket (Majiduddin et al., 2002).

**Figure 5.**
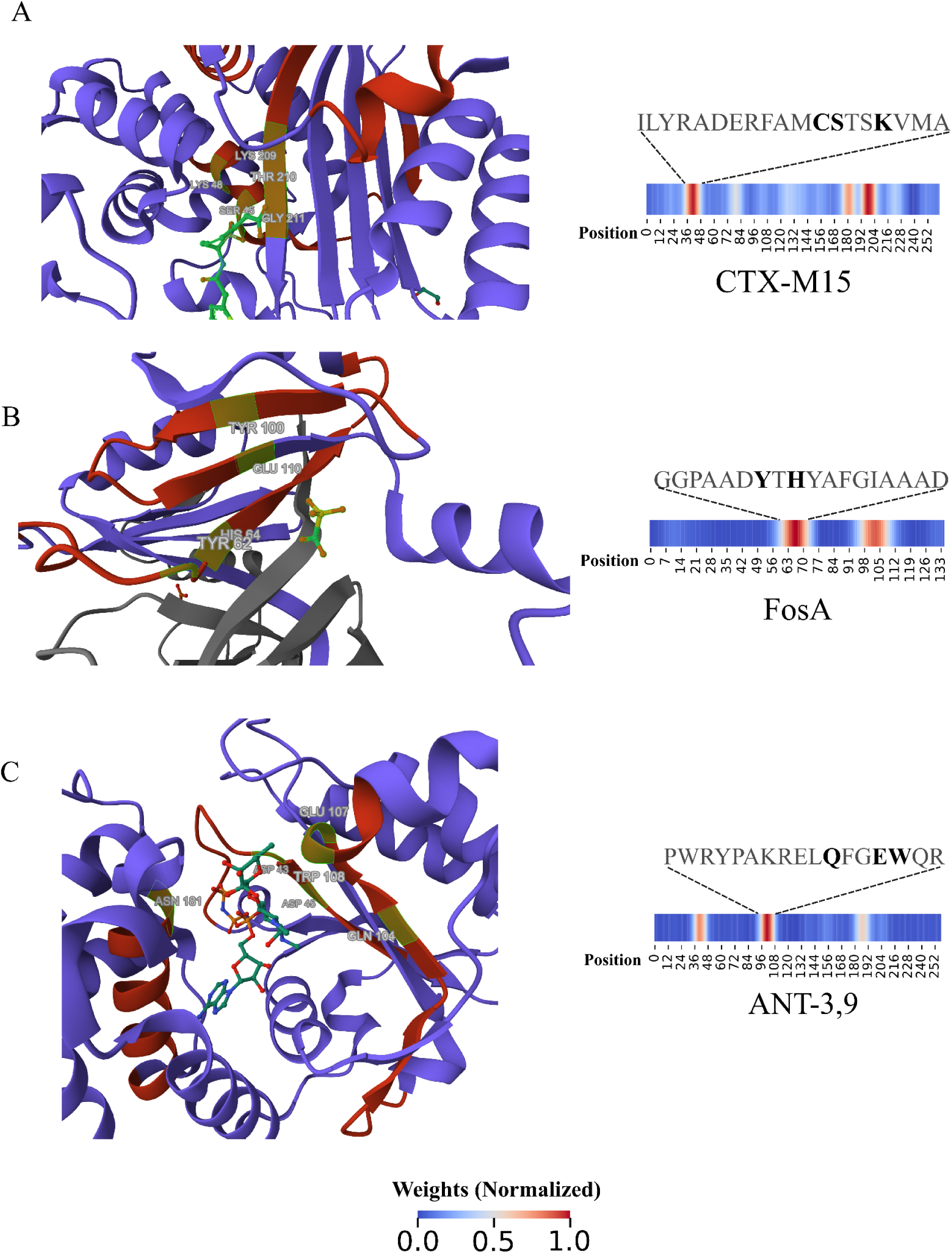
CNN model interpretability. PDB 3D structures of a CTX-M beta-lactamase (A), a fosfomycin thiol transferase (B), and an aminoglycoside nucleotidyltransferase (C) colored according to CNN neuron firing patterns. The green-highlighted residues on the structures and the black-highlighted above heatmaps are annotated as binding sites to antimicrobial molecules. The grey structure on B is the other fosfomycin thiol transferase monomer.

**Table 3.**
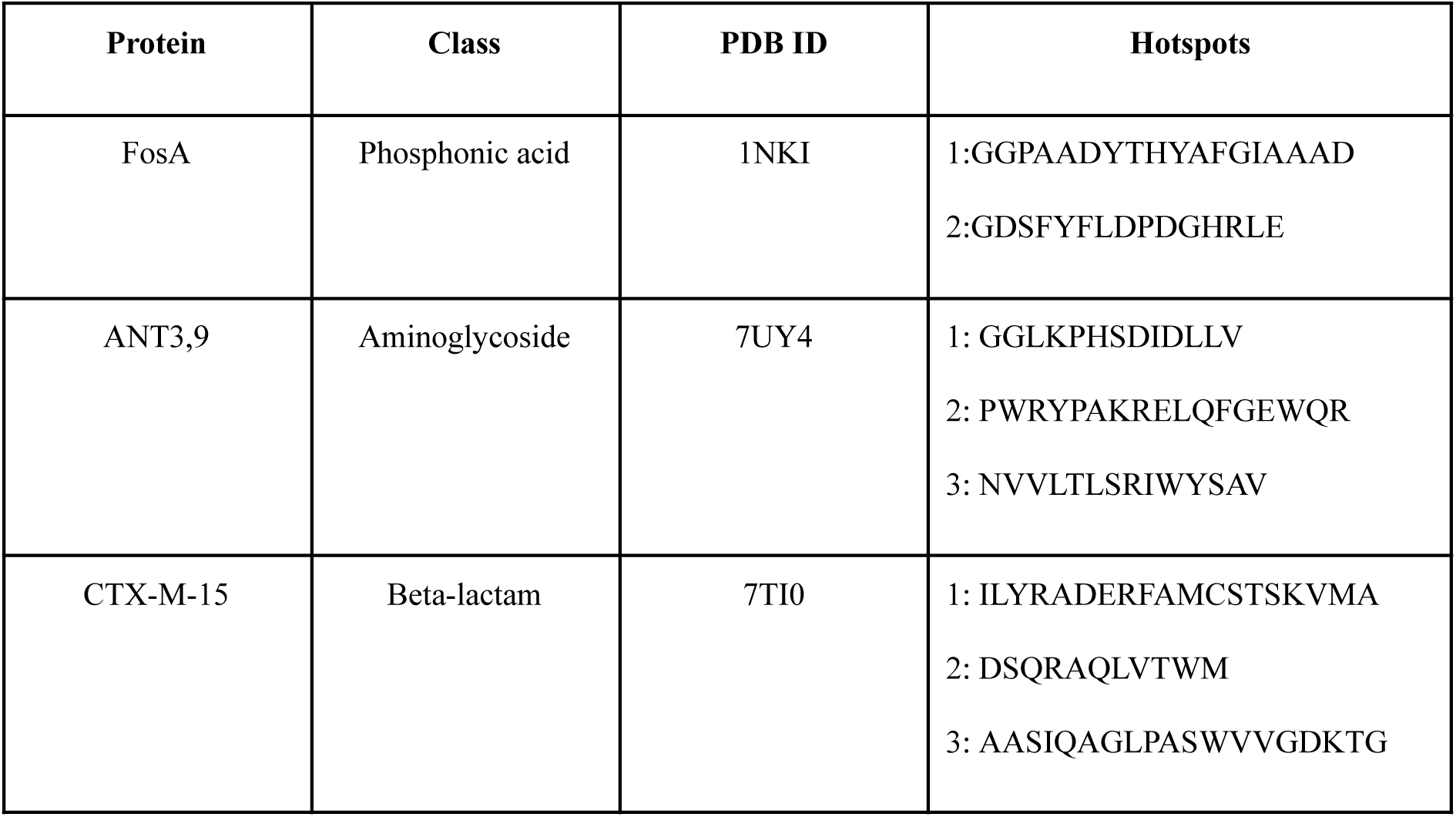
Proteins from PDB and subsequences of high importance for classification.

For the FosA (PDB: 1NKI) protein (Figure 5B), our model highlighted two hotspots that build the catalyst site where the residues H64, Y100, and E110 bind the fosfomycin molecule and Y62, which bridges the active site and the dimer interface loop (Klontz et al., 2017). Similarly, the aminoglycoside (3’’) (9) adenylyltransferase (PDB: 7UY4) (Figure 5C) presented residues D43, D45, Q104, E107, W108, and D181 annotated as binding sites for spectinomycin antimicrobial (Kanchugal P and Selmer, 2020). These results confirmed that our model learned biological features from proteins without reference sequences or sequence alignment comparisons.

We checked our model misclassifications and found evidence of misannotated proteins in the NCRD95 dataset that were correctly classified by our CNN. In our test, the protein NR_MCR0138297.1 is labeled as aminoglycoside resistance and was classified as beta-lactam. BLASTP revealed that NR_MCR0138297.1 is 86.6% identical to a PAC-1 (NR_ABP88743.1) beta-lactamase in our training set. The same was observed for NR_WP_193096997.1, a member of the aminoglycoside resistance class that is 52.1% similar to Arr-1 (NR_PRC48666.1), a rifamycin-resistant protein.

A third example of misannotation from NCRD95 is the oleD protein, a glycosyltransferase that confers resistance only to macrolide antimicrobials (Bolam et al., 2007; Quirós et al., 2000). In this case, the protein NR_WP_166002910.1 in the training set and the NR_PZH16762.1 protein in the test set were both labeled as MLS resistance class and shared 82.8% of protein identity. However, our CNN classified NR_WP_166002910.1 as a macrolide-resistant protein.

### DeepSEA tool

To make our model available for the scientific community, the CNN was wrapped in a command-line interface tool available on https://github.com/computational-chemical-biology/DeepSEA-project. There are two usage options for DeepSEA, for resistance classification the user can provide a raw FASTA file as input and obtain a table format file with model predictions and their respective probabilities, or for motif detection, a multisequence alignment FASTA file can be analyzed by DeepSEA, returning a heatmap with the hotspots for each aligned sequence.

## Discussion

In this work, we developed an alignment-free tool that functionally annotates proteins across multiple antimicrobial resistance classes and distinguishes resistance from nonresistance proteins without protein alignment. The DeepSEA model was trained to recognize motifs (hotspots) from resistance proteins and therefore provides an explanation for its predictions, which helps users from biological backgrounds to interpret the results. Benchmarking revealed that DeepSEA is superior to protein alignment when it comes to identifying fewer false negatives.

The protein alignment model uses homology inference (Pearson, 2013) as a shortcut from sequence to function in the classical sequence-structure-function paradigm (Koehler Leman et al., 2023). However, even homologous groups, such as serine beta-lactamases, can vary considerably outside their active sites (Joris et al., 1988b), forcing the databases to have reference sequences for variants, otherwise leading to high rates of false negative results (Arango-Argoty et al., 2018).

The lower recall values of RGI and AMRFinder demonstrated that the lack of proper reference sequences in their databases led to several misclassifications of resistant proteins as non-resistant proteins. For instance, RGI and ARMFinder classified 88% and 79% of the proteins that resist glycopeptides as non-resistant proteins, respectively. Moreover, both alignment-based tools presented misleading results for proteins related to resistance to beta-lactam antimicrobials. DeepSEA, on the other hand, had only 8 beta-lactam resistance proteins misclassified as nonresistance.

DeepSEA performance matched ESM2 even though the ESM2 model is composed of a transformer-based architecture pre-trained on the approximately 60 million proteins from the UniRef database (September 2021 version). Moreover, the CNN-based architecture allowed DeepSEA’s output interpretability without data perturbation. For example, to explain HMD-ARG (Li et al., 2021) results, the author had to investigate the effect of *in silico* mutations in protein sites on model performance. This need for checking protein variations is computationally expensive and infeasible in most cases. The pioneer DeepARG (Arango-Argoty et al., 2018) used a multilayer perceptron model, which can not offer explainability, and also needs to prefilter resistance-like proteins via DIAMOND (Buchfink et al., 2015) before model prediction itself.

During the benchmarking experiments, the DeepSEA precision value was lower for nonresistance than for other classes. This was expected due to the miscellaneous composition of the nonresistance class, which contains proteins with a plethora of functions. This lack of homogeneity of the nonresistance class forces the CNN neurons to fire at patterns restricted to the other classes. We extracted this information directly from the model and converted it to a human-readable format, contextualizing DeepSEA’s output to information provided by multisequence alignment. This feature could be used in future research to combine 3D structure prediction with functional annotations.

DeepSEA was designed to address a key issue in antimicrobial resistance protein annotation: the false negative rates. We focused on pushing the frontiers of protein annotation made by sequence similarity. Therefore, DeepSEA does not contain genome assemblers and open reading frame predictors. Also, since genomic research pipelines often use annotation tools of a general purpose such as Prokka (Seemann, 2014), DeepSEA can be employed to reannotate proteins previously assigned as hypothetical and increase the knowledge on the diversity of proteins produced by bacteria or to screen large datasets of unannotated protein sequences predicted by other tools. DeepSEA was designed to find signatures of proteins for broad resistance classes. For instance, future work will address more specific questions and split the beta-lactam class according to the type of beta-lactam antimicrobial (i.e., cephalosporin, penem, to cite a few).

## Conclusion

In this work, we addressed the high rates of false negatives in antimicrobial resistance protein annotation by sequence alignment. It was demonstrated that a single CNN model, trained directly on protein sequences instead of MSA, achieved higher performance than alignment-based methods and produced only a few false negatives for every protein class in the training set without detectable overfitting. Moreover, we proposed an algorithm to extract and transform information from the CNN into meaningful human-readable insights about the model’s decision-making that help users from different backgrounds interpret the results. Future works could use multimodal data to expand these explanatory features from proteins to complex metabolic pathways of resistant bacteria to reveal protein-protein interactions that contribute to the resistant phenotype.

## List of abbreviations

MSA: Multiple Sequence Alignment
CNN: Convolutional Neural Network
RGI: Resistance Gene Identifiers
NCRD: Non-redundant Comprehensive Resistance Database
AMR: Antimicrobial Resistance
HIV: Human Immunodeficiency Virus
AST: Antimicrobial Susceptibility Tests
PDB: Protein Data Bank
EMS: Evolutionary Scale Model
NonR: Non-resistant proteins
MLS: Macrolide, lincosamide, and streptogramin

## Declarations

### Ethics approval and consent to participate

Not applicable

### Consent for publication

Not applicable

### Availability of data and material

DeepSEA datasets and model can be accessed and downloaded from the project’s repository on GitHub: https://github.com/computational-chemical-biology/DeepSEA-project.

### Authors’ contributions

TCB: data collection and curation, model training and evaluation, data analysis, and manuscript writing.

ARP: manuscript review and data analysis review.

RRS: model evaluation, data analysis review, manuscript writing, and manuscript review.

## Acknowledgments

We are grateful to MSc Guilherme Marcelino Viana de Siqueira for his valuable and insightful ideas that enriched the conceptual framework of this study.

## Bibliography

Alcock, B.P., Raphenya, A.R., Lau, T.T.Y., Tsang, K.K., Bouchard, M., Edalatmand, A., Huynh, W., Nguyen, A.-L.V., Cheng, A.A., Liu, S., Min, S.Y., Miroshnichenko, A., Tran, H.-K., Werfalli, R.E., Nasir, J.A., Oloni, M., Speicher, D.J., Florescu, A., Singh, B., Faltyn, M., Hernandez-Koutoucheva, A., Sharma, A.N., Bordeleau, E., Pawlowski, A.C., Zubyk, H.L., Dooley, D., Griffiths, E., Maguire, F., Winsor, G.L., Beiko, R.G., Brinkman, F.S.L., Hsiao, W.W.L., Domselaar, G.V., McArthur, A.G., 2020. CARD 2020: antibiotic resistome surveillance with the comprehensive antibiotic resistance database. Nucleic Acids Res. 48, D517–D525. 10.1093/nar/gkz935

Arango-Argoty, G., Garner, E., Pruden, A., Heath, L.S., Vikesland, P., Zhang, L., 2018. DeepARG: a deep learning approach for predicting antibiotic resistance genes from metagenomic data. Microbiome 6, 23. 10.1186/s40168-018-0401-z

Bernett, J., Blumenthal, D.B., Grimm, D.G., Haselbeck, F., Joeres, R., Kalinina, O.V., List, M., 2024. Guiding questions to avoid data leakage in biological machine learning applications. Nat. Methods 21, 1444–1453. 10.1038/s41592-024-02362-y

Bileschi, M.L., Belanger, D., Bryant, D.H., Sanderson, T., Carter, B., Sculley, D., Bateman, A., DePristo, M.A., Colwell, L.J., 2022. Using deep learning to annotate the protein universe. Nat. Biotechnol. 40, 932–937. 10.1038/s41587-021-01179-w

Biswal, S., Caetano, K., Jain, D., Sarrila, A., Munshi, T., Dickman, R., Tabor, A.B., Rath, S.N., Bhakta, S., Ghosh, A.S., 2023. Antimicrobial Peptides Designed against the Ω-Loop of Class A β-Lactamases to Potentiate the Efficacy of β-Lactam Antibiotics. Antibiotics 12, 553. 10.3390/antibiotics12030553

Blum, M., Chang, H.-Y., Chuguransky, S., Grego, T., Kandasaamy, S., Mitchell, A., Nuka, G., Paysan-Lafosse, T., Qureshi, M., Raj, S., Richardson, L., Salazar, G.A., Williams, L., Bork, P., Bridge, A., Gough, J., Haft, D.H., Letunic, I., Marchler-Bauer, A., Mi, H., Natale, D.A., Necci, M., Orengo, C.A., Pandurangan, A.P., Rivoire, C., Sigrist, C.J.A., Sillitoe, I., Thanki, N., Thomas, P.D., Tosatto, S.C.E., Wu, C.H., Bateman, A., Finn, R.D., 2021. The InterPro protein families and domains database: 20 years on. Nucleic Acids Res. 49, D344–D354. 10.1093/nar/gkaa977

Bolam, D.N., Roberts, S., Proctor, M.R., Turkenburg, J.P., Dodson, E.J., Martinez-Fleites, C., Yang, M., Davis, B.G., Davies, G.J., Gilbert, H.J., 2007. The crystal structure of two macrolide glycosyltransferases provides a blueprint for host cell antibiotic immunity. Proc. Natl. Acad. Sci. U. S. A. 104, 5336–5341. 10.1073/pnas.0607897104

Buchfink, B., Xie, C., Huson, D.H., 2015. Fast and sensitive protein alignment using DIAMOND. Nat. Methods 12, 59–60. 10.1038/nmeth.3176

Djordjevic, S.P., Jarocki, V.M., Seemann, T., Cummins, M.L., Watt, A.E., Drigo, B., Wyrsch, E.R., Reid, C.J., Donner, E., Howden, B.P., 2023. Genomic surveillance for antimicrobial resistance — a One Health perspective. Nat. Rev. Genet. 1–16. 10.1038/s41576-023-00649-y

Feldgarden, M., Brover, V., Haft, D.H., Prasad, A.B., Slotta, D.J., Tolstoy, I., Tyson, G.H., Zhao, S., Hsu, C.-H., McDermott, P.F., Tadesse, D.A., Morales, C., Simmons, M., Tillman, G., Wasilenko, J., Folster, J.P., Klimke, W., 2019. Validating the AMRFinder Tool and Resistance Gene Database by Using Antimicrobial Resistance Genotype-Phenotype Correlations in a Collection of Isolates. Antimicrob. Agents Chemother. 63, e00483-19. 10.1128/AAC.00483-19

Fleming, A., 1929. On the Antibacterial Action of Cultures of a Penicillium, with Special Reference to their Use in the Isolation of B. influenzæ. Br. J. Exp. Pathol. 10, 226–236.

Florensa, A.F., Kaas, R.S., Clausen, P.T.L.C., Aytan-Aktug, D., Aarestrup, F.M., 2022. ResFinder – an open online resource for identification of antimicrobial resistance genes in next-generation sequencing data and prediction of phenotypes from genotypes. Microb. Genomics 8, 000748. 10.1099/mgen.0.000748

Ghuysen, J.-M., 1991. Serine β-lactamases and penicillin-binding proteins. Annu. Rev. Microbiol. 45. 10.1146/annurev.mi.45.100191.000345

Hasman, H., Saputra, D., Sicheritz-Ponten, T., Lund, O., Svendsen, C.A., Frimodt-Møller, N., Aarestrup, F.M., 2014. Rapid Whole-Genome Sequencing for Detection and Characterization of Microorganisms Directly from Clinical Samples. J. Clin. Microbiol. 52, 139–146. 10.1128/JCM.02452-13

Jones, P., Binns, D., Chang, H.-Y., Fraser, M., Li, W., McAnulla, C., McWilliam, H., Maslen, J., Mitchell, A., Nuka, G., Pesseat, S., Quinn, A.F., Sangrador-Vegas, A., Scheremetjew, M., Yong, S.-Y., Lopez, R., Hunter, S., 2014. InterProScan 5: genome-scale protein function classification. Bioinforma. Oxf. Engl. 30, 1236–1240. 10.1093/bioinformatics/btu031

Joris, B., Ghuysen, J.M., Dive, G., Renard, A., Dideberg, O., Charlier, P., Frère, J.M., Kelly, J.A., Boyington, J.C., Moews, P.C., 1988a. The active-site-serine penicillin-recognizing enzymes as members of the Streptomyces R61 DD-peptidase family. Biochem. J. 250, 313–324.

Joris, B., Ghuysen, J.M., Dive, G., Renard, A., Dideberg, O., Charlier, P., Frère, J.M., Kelly, J.A., Boyington, J.C., Moews, P.C., 1988b. The active-site-serine penicillin-recognizing enzymes as members of the Streptomyces R61 DD-peptidase family. Biochem. J. 250, 313–324.

Judge, A., Hu, L., Sankaran, B., Van Riper, J., Venkataram Prasad, B.V., Palzkill, T., 2023. Mapping the determinants of catalysis and substrate specificity of the antibiotic resistance enzyme CTX-M β-lactamase. Commun. Biol. 6, 1–11. 10.1038/s42003-023-04422-z

Kanchugal P, S., Selmer, M., 2020. Structural Recognition of Spectinomycin by Resistance Enzyme ANT(9) from Enterococcus faecalis. Antimicrob. Agents Chemother. 64, 10.1128/aac.00371-20. 10.1128/aac.00371-20

Klontz, E.H., Tomich, A.D., Günther, S., Lemkul, J.A., Deredge, D., Silverstein, Z., Shaw, J.F., McElheny, C., Doi, Y., Wintrode, P.L., MacKerell, A.D., Sluis-Cremer, N., Sundberg, E.J., 2017. Structure and Dynamics of FosA-Mediated Fosfomycin Resistance in Klebsiella pneumoniae and Escherichia coli. Antimicrob. Agents Chemother. 61, 10.1128/aac.01572-17. 10.1128/aac.01572-17

Koehler Leman, J., Szczerbiak, P., Renfrew, P.D., Gligorijevic, V., Berenberg, D., Vatanen, T., Taylor, B.C., Chandler, C., Janssen, S., Pataki, A., Carriero, N., Fisk, I., Xavier, R.J., Knight, R., Bonneau, R., Kosciolek, T., 2023. Sequence-structure-function relationships in the microbial protein universe. Nat. Commun. 14, 2351. 10.1038/s41467-023-37896-w

Leigh Van Valen, 1973. A New Evolutionary Law. NEW Evol. LAW.

Li, W., Godzik, A., 2006. Cd-hit: a fast program for clustering and comparing large sets of protein or nucleotide sequences. Bioinforma. Oxf. Engl. 22, 1658–1659. 10.1093/bioinformatics/btl158

Li, Y., Xu, Z., Han, W., Cao, H., Umarov, R., Yan, A., Fan, M., Chen, H., Duarte, C.M., Li, L., Ho, P.-L., Gao, X., 2021. HMD-ARG: hierarchical multi-task deep learning for annotating antibiotic resistance genes. Microbiome 9, 40. 10.1186/s40168-021-01002-3

Lin, Z., Akin, H., Rao, R., Hie, B., Zhu, Z., Lu, W., Smetanin, N., Verkuil, R., Kabeli, O., Shmueli, Y., dos Santos Costa, A., Fazel-Zarandi, M., Sercu, T., Candido, S., Rives, A., 2023. Evolutionary-scale prediction of atomic-level protein structure with a language model. Science 379, 1123–1130. 10.1126/science.ade2574

Liu, B., Pop, M., 2009. ARDB—Antibiotic Resistance Genes Database. Nucleic Acids Res. 37, D443–D447. 10.1093/nar/gkn656

Majiduddin, F.K., Materon, I.C., Palzkill, T.G., 2002. Molecular analysis of beta-lactamase structure and function. Int. J. Med. Microbiol. 292, 127–137. 10.1078/1438-4221-00198

Mao, Y., Liu, X., Zhang, N., Wang, Z., Han, M., 2023a. NCRD: A non-redundant comprehensive database for detecting antibiotic resistance genes. iScience 26, 108141. 10.1016/j.isci.2023.108141

Mao, Y., Liu, X., Zhang, N., Wang, Z., Han, M., 2023b. NCRD: A non-redundant comprehensive database for detecting antibiotic resistance genes. iScience 26, 108141. 10.1016/j.isci.2023.108141

Murray, C.J.L., Ikuta, K.S., Sharara, F., Swetschinski, L., Aguilar, G.R., Gray, A., Han, C., Bisignano, C., Rao, P., Wool, E., Johnson, S.C., Browne, A.J., Chipeta, M.G., Fell, F., Hackett, S., Haines-Woodhouse, G., Hamadani, B.H.K., Kumaran, E.A.P., McManigal, B., Achalapong, S., Agarwal, R., Akech, S., Albertson, S., Amuasi, J., Andrews, J., Aravkin, A., Ashley, E., Babin, F.-X., Bailey, F., Baker, S., Basnyat, B., Bekker, A., Bender, R., Berkley, J.A., Bethou, A., Bielicki, J., Boonkasidecha, S., Bukosia, J., Carvalheiro, C., Castañeda-Orjuela, C., Chansamouth, V., Chaurasia, S., Chiurchiù, S., Chowdhury, F., Donatien, R.C., Cook, A.J., Cooper, B., Cressey, T.R., Criollo-Mora, E., Cunningham, M., Darboe, S., Day, N.P.J., Luca, M.D., Dokova, K., Dramowski, A., Dunachie, S.J., Bich, T.D., Eckmanns, T., Eibach, D., Emami, A., Feasey, N., Fisher-Pearson, N., Forrest, K., Garcia, C., Garrett, D., Gastmeier, P., Giref, A.Z., Greer, R.C., Gupta, V., Haller, S., Haselbeck, A., Hay, S.I., Holm, M., Hopkins, S., Hsia, Y., Iregbu, K.C., Jacobs, J., Jarovsky, D., Javanmardi, F., Jenney, A.W.J., Khorana, M., Khusuwan, S., Kissoon, N., Kobeissi, E., Kostyanev, T., Krapp, F., Krumkamp, R., Kumar, A., Kyu, H.H., Lim, C., Lim, K., Limmathurotsakul, D., Loftus, M.J., Lunn, M., Ma, J., Manoharan, A., Marks, F., May, J., Mayxay, M., Mturi, N., Munera-Huertas, T., Musicha, P., Musila, L.A., Mussi-Pinhata, M.M., Naidu, R.N., Nakamura, T., Nanavati, R., Nangia, S., Newton, P., Ngoun, C., Novotney, A., Nwakanma, D., Obiero, C.W., Ochoa, T.J., Olivas-Martinez, A., Olliaro, P., Ooko, E., Ortiz-Brizuela, E., Ounchanum, P., Pak, G.D., Paredes, J.L., Peleg, A.Y., Perrone, C., Phe, T., Phommasone, K., Plakkal, N., Ponce-de-Leon, A., Raad, M., Ramdin, T., Rattanavong, S., Riddell, A., Roberts, T., Robotham, J.V., Roca, A., Rosenthal, V.D., Rudd, K.E., Russell, N., Sader, H.S., Saengchan, W., Schnall, J., Scott, J.A.G., Seekaew, S., Sharland, M., Shivamallappa, M., Sifuentes-Osornio, J., Simpson, A.J., Steenkeste, N., Stewardson, A.J., Stoeva, T., Tasak, N., Thaiprakong, A., Thwaites, G., Tigoi, C., Turner, C., Turner, P., Doorn, H.R. van, Velaphi, S., Vongpradith, A., Vongsouvath, M., Vu, H., Walsh, T., Walson, J.L., Waner, S., Wangrangsimakul, T., Wannapinij, P., Wozniak, T., Sharma, T.E.M.W.Y., Yu, K.C., Zheng, P., Sartorius, B., Lopez, A.D., Stergachis, A., Moore, C., Dolecek, C., Naghavi, M., 2022. Global burden of bacterial antimicrobial resistance in 2019: a systematic analysis. The Lancet 399, 629–655. 10.1016/S0140-6736(21)02724-0

Pal, C., Bengtsson-Palme, J., Kristiansson, E., Larsson, D.G.J., 2016. The structure and diversity of human, animal and environmental resistomes. Microbiome 4, 54. 10.1186/s40168-016-0199-5

Paszke, A., Gross, S., Massa, F., Lerer, A., Bradbury, J., Chanan, G., Killeen, T., Lin, Z., Gimelshein, N., Antiga, L., Desmaison, A., Köpf, A., Yang, E., DeVito, Z., Raison, M., Tejani, A., Chilamkurthy, S., Steiner, B., Fang, L., Bai, J., Chintala, S., 2019. PyTorch: an imperative style, high-performance deep learning library, in: Proceedings of the 33rd International Conference on Neural Information Processing Systems. Curran Associates Inc., Red Hook, NY, USA, pp. 8026–8037.

Pearson, W.R., 2013. An Introduction to Sequence Similarity (“Homology”) Searching. Curr. Protoc. Bioinforma. Ed. Board Andreas Baxevanis Al 0 3, 10.1002/0471250953.bi0301s42. 10.1002/0471250953.bi0301s42

Quirós, L.M., Carbajo, R.J., Braña, A.F., Salas, J.A., 2000. Glycosylation of Macrolide Antibiotics: PURIFICATION AND KINETIC STUDIES OF A MACROLIDE GLYCOSYLTRANSFERASE FROM STREPTOMYCES ANTIBIOTICUS*. J. Biol. Chem. 275, 11713–11720. 10.1074/jbc.275.16.11713

Seemann, T., 2014. Prokka: rapid prokaryotic genome annotation. Bioinformatics 30, 2068–2069. 10.1093/bioinformatics/btu153

Shen, W., Sipos, B., Zhao, L., 2024. SeqKit2: A Swiss army knife for sequence and alignment processing. iMeta 3, e191. 10.1002/imt2.191

Skinnider, M.A., Johnston, C.W., Gunabalasingam, M., Merwin, N.J., Kieliszek, A.M., MacLellan, R.J., Li, H., Ranieri, M.R.M., Webster, A.L.H., Cao, M.P.T., Pfeifle, A., Spencer, N., To, Q.H., Wallace, D.P., Dejong, C.A., Magarvey, N.A., 2020. Comprehensive prediction of secondary metabolite structure and biological activity from microbial genome sequences. Nat. Commun. 11, 6058. 10.1038/s41467-020-19986-1

ur Rahman, S., Ali, T., Ali, I., Khan, N.A., Han, B., Gao, J., 2018. The Growing Genetic and Functional Diversity of Extended Spectrum Beta-Lactamases. BioMed Res. Int. 2018, 9519718. 10.1155/2018/9519718

Wang, B., Mei, C., Wang, Y., Zhou, Y., Cheng, M.-T., Zheng, C.-H., Wang, L., Zhang, J., Chen, P., Xiong, Y., 2021. Imbalance Data Processing Strategy for Protein Interaction Sites Prediction. IEEE/ACM Trans. Comput. Biol. Bioinform. 18, 985–994. 10.1109/TCBB.2019.2953908

Wang, Z., Li, S., You, R., Zhu, S., Zhou, X.J., Sun, F., 2021. ARG-SHINE: improve antibiotic resistance class prediction by integrating sequence homology, functional information and deep convolutional neural network. NAR Genomics Bioinforma. 3, lqab066. 10.1093/nargab/lqab066

Wolf, T., Debut, L., Sanh, V., Chaumond, J., Delangue, C., Moi, A., Cistac, P., Rault, T., Louf, R., Funtowicz, M., Davison, J., Shleifer, S., von Platen, P., Ma, C., Jernite, Y., Plu, J., Xu, C., Le Scao, T., Gugger, S., Drame, M., Lhoest, Q., Rush, A., 2020. Transformers: State-of-the-Art Natural Language Processing, in: Liu, Q., Schlangen, D. (Eds.), Proceedings of the 2020 Conference on Empirical Methods in Natural Language Processing: System Demonstrations. Association for Computational Linguistics, Online, pp. 38–45. 10.18653/v1/2020.emnlp-demos.6

Wu, J., Ouyang, J., Qin, H., Zhou, J., Roberts, R., Siam, R., Wang, L., Tong, W., Liu, Z., Shi, T., 2023. PLM-ARG: antibiotic resistance gene identification using a pretrained protein language model. Bioinformatics 39, btad690. 10.1093/bioinformatics/btad690

Yin, X., Jiang, X.-T., Chai, B., Li, L., Yang, Y., Cole, J.R., Tiedje, J.M., Zhang, T., 2018. ARGs-OAP v2.0 with an expanded SARG database and Hidden Markov Models for enhancement characterization and quantification of antibiotic resistance genes in environmental metagenomes. Bioinformatics 34, 2263–2270. 10.1093/bioinformatics/bty053

Yoshikawa, T.T., 2002. Antimicrobial Resistance and Aging: Beginning of the End of the Antibiotic Era? J. Am. Geriatr. Soc. 50, 226–229. 10.1046/j.1532-5415.50.7s.2.x

