## Additional File 2 for "DeepSEA: an alignment-free explainable approach to annotate antimicrobial resistance proteins"

Supplementary Table 1. Class weights and proportions

| Class | Proportion in the dataset (%) | Weight |
|---|---|---|
| MLS | 3 | 3.03 |
| NonR | 26 | 0.39 |
| Aminoglycoside | 7 | 1.40 |
| beta-lactam | 11 | 0.91 |
| chloramphenicol | 3 | 3.96 |
| glycopeptide | 35 | 0.28 |
| macrolide | 1 | 9.98 |
| phosphonic acid | 4 | 2.44 |
| rifamycin | 3 | 2.94 |
| tetracycline | 6 | 1.57 |

Supplementary Table 2. CNN detailed architecture.

| **Layer** | **Hyperparameters** |
|---|---|
| TextVectorization | max tokens = 20<br>output_sequence_length = 1024 |
| Embedding | input dim = 20<br>output dim = 50 |
| Conv1D | filters (kernels) = 968<br>kernel size = 9<br>activation = relu<br>padding = same |
| Dropout | rate = 0.25<br>seed = 42 |
| Conv1D | filters (kernels) = 464<br>kernel size = 9<br>activation = relu<br>padding = same |
| Dropout | rate = 0.25 |

| Layer | Hyperparameters |
|---|---|
| | seed = 42 |
| Conv1D | filters (kernels) = 296<br>kernel size = 9<br>activation = relu<br>padding = same |
| Dropout | rate = 0.25<br>seed = 42 |
| Conv1D | filters (kernels) = 520<br>kernel size = 9<br>activation = relu<br>padding = same |
| GlobalAveragePooling1D | - |
| Dense | neurons = 10<br>activation = softmax |
