## Additional File 1 for "DeepSEA: an alignment-free explainable approach to annotate antimicrobial resistance proteins"

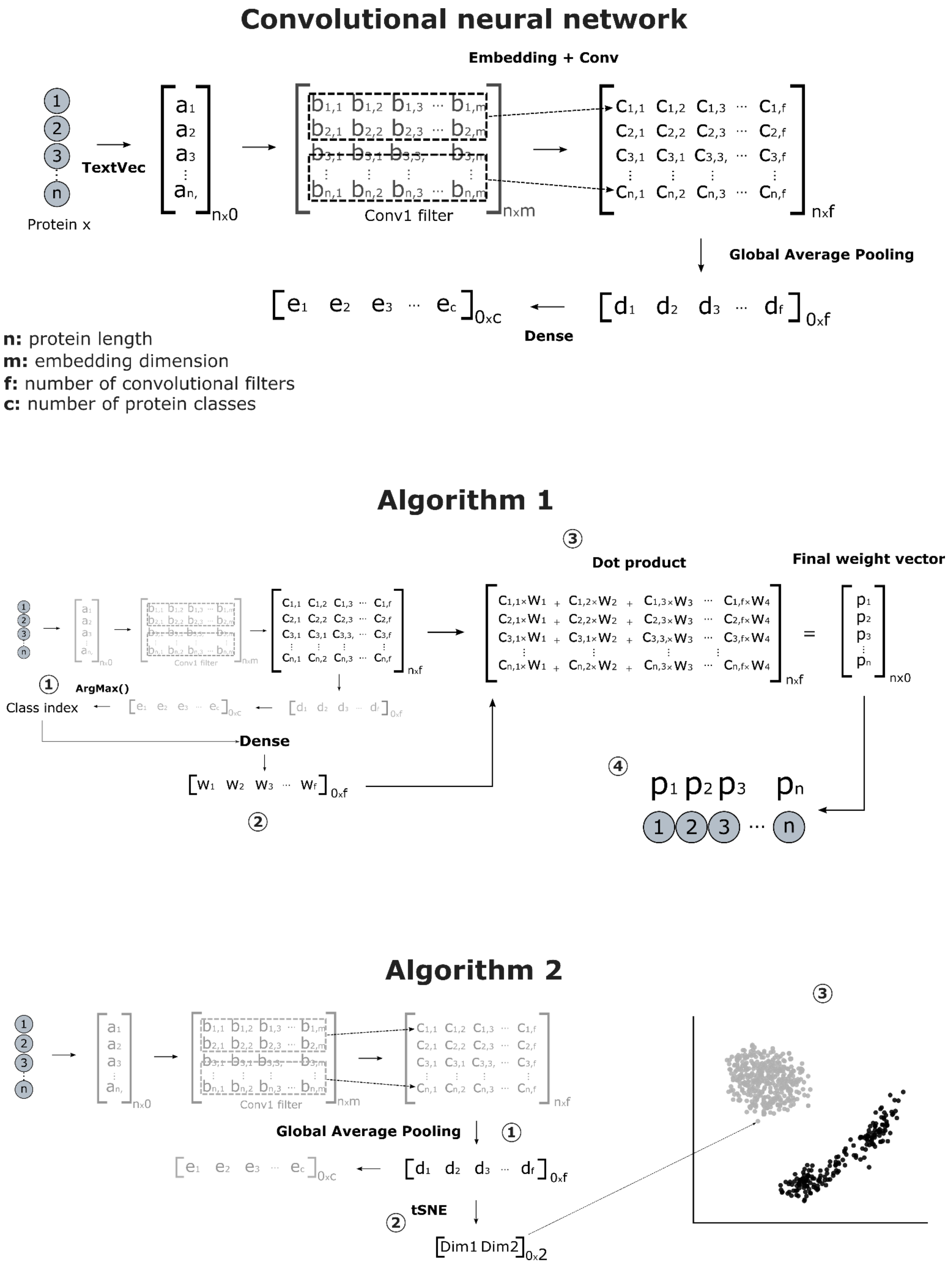


Supplementary Figure 1. Visual representation of information workflow inside a convolutional neural network (CNN) and mathematical transformation of CNN internal states by Algorithms 1 and 2 into a human-interpretable output.

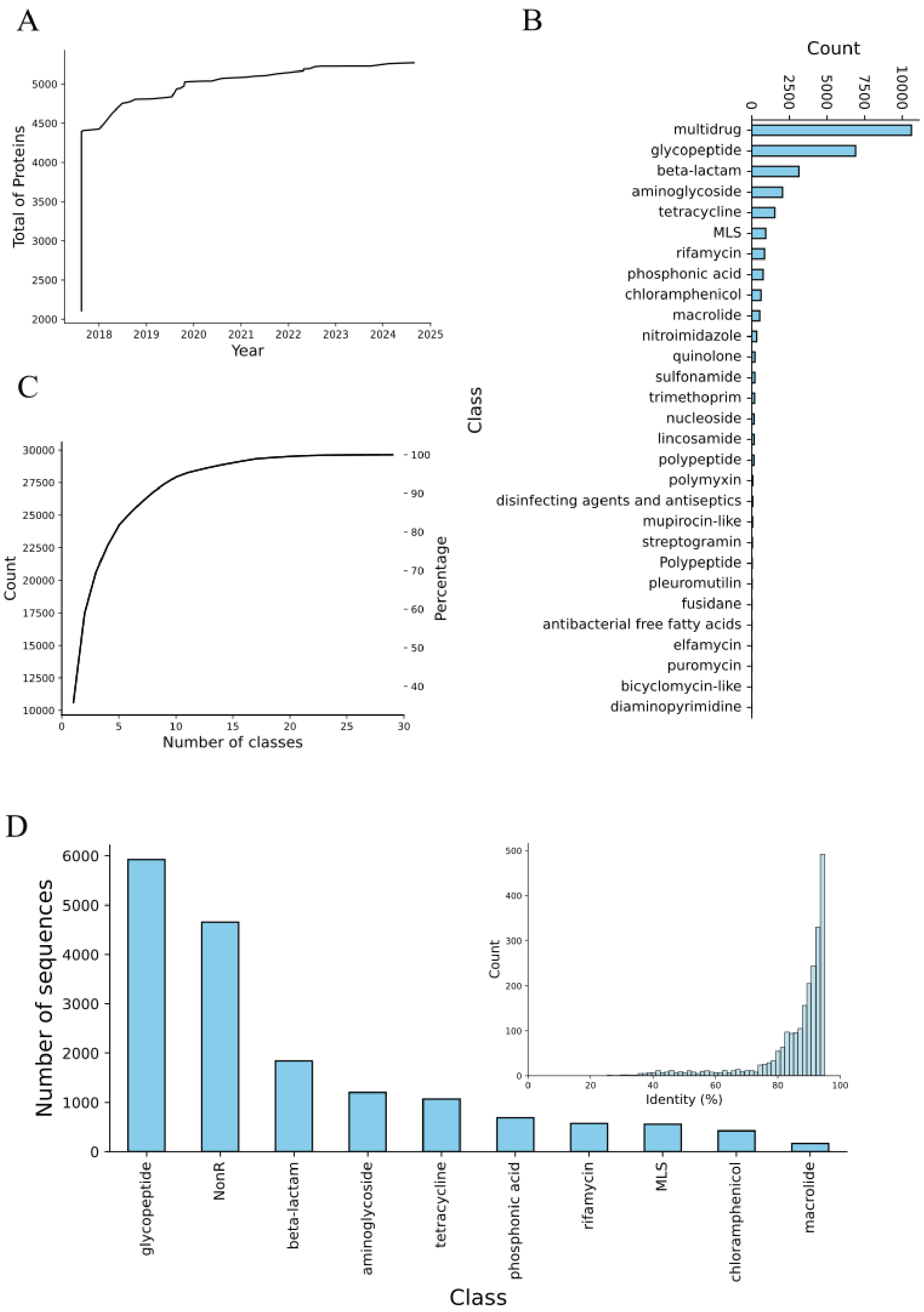


Supplementary Figure 2. Database description. A) Cumulative curve of CARD database updates from early 2017 to late 2024. C) Cumulative curve of the original antimicrobial resistance protein classes. D) The distribution of classes for CNN development after selecting the most abundant ones. D) Final class distribution and protein similarity between training and test sets. The alignment was performed considering the converge parameter equals 100% to avoid local alignment and e-value < 0.001.

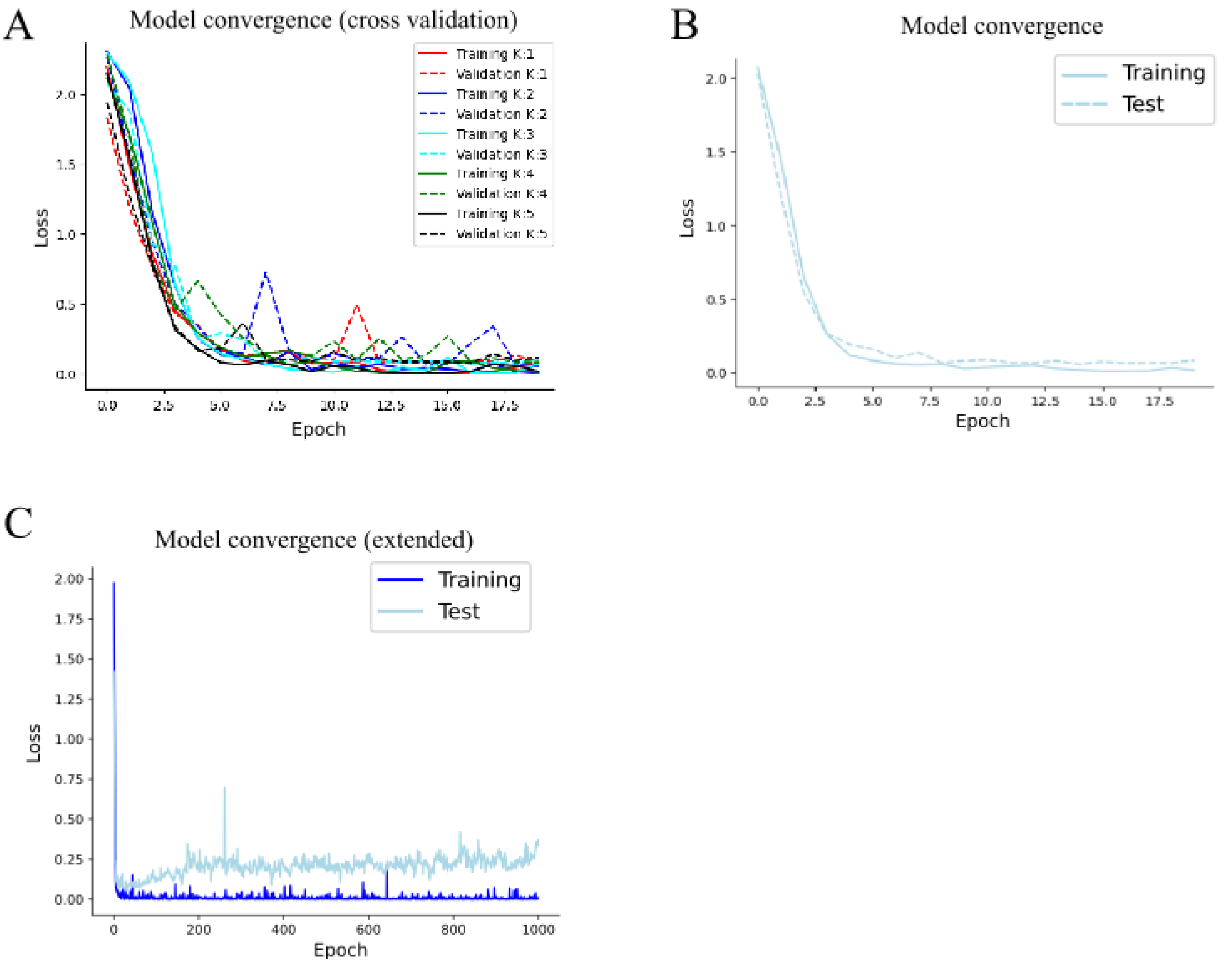


Supplementary Figure 3. Model convergences. A) Convergence curve for the CNN model trained using the complete training set. The limited number of epochs is due to early stopping to avoid overfitting. The dashed curve represented the loss calculator using the holdout test set. B) Convergence curve for an extended period (1000 epochs) for checking overfitting. C) Converge curves from 5-fold cross-validation. The training set was randomly subdivided at five different points as detailed in Methods.

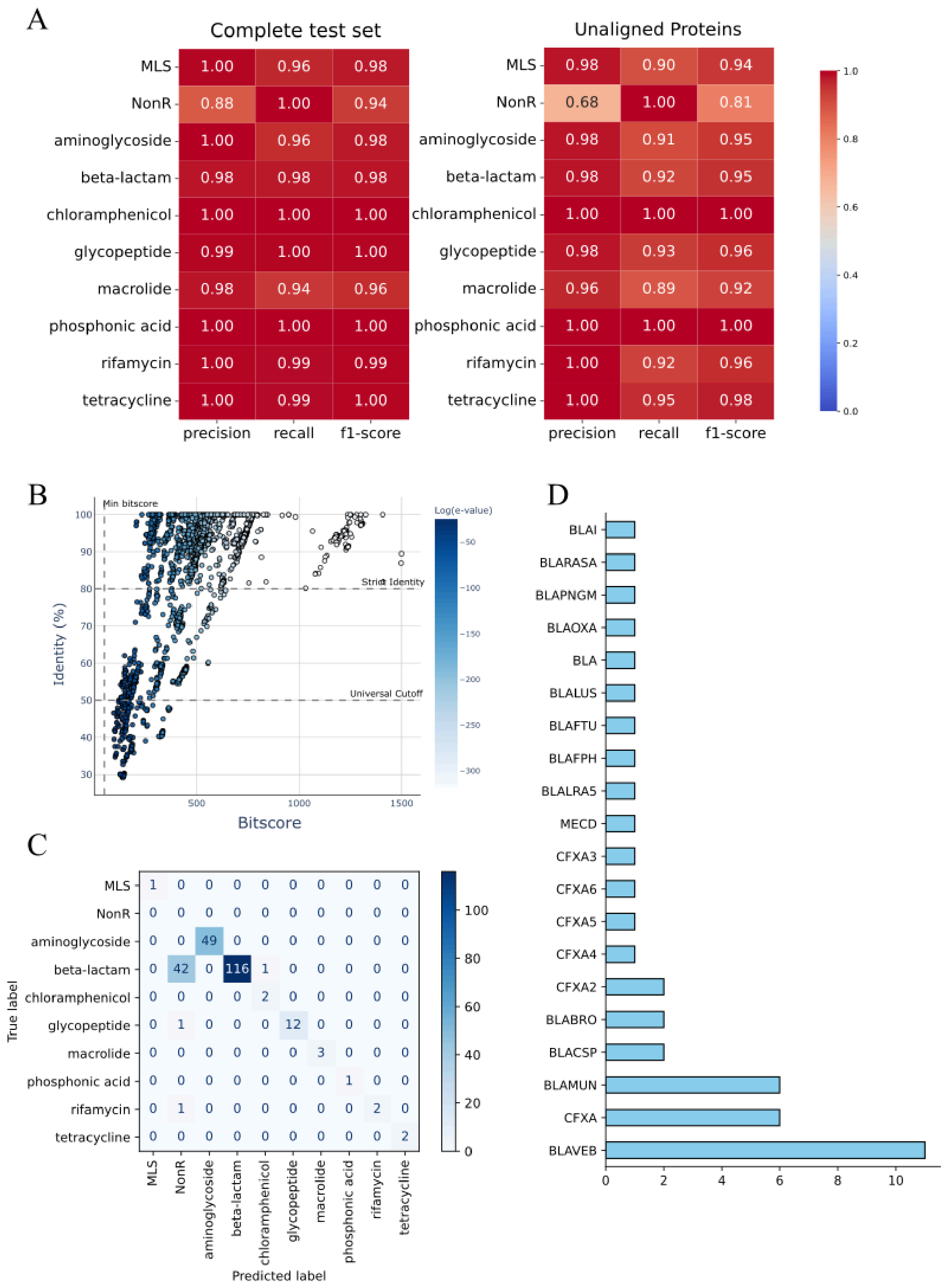


Supplementary Figure 4. The limits of generalization. A) Evaluation metrics of the entire test set (n=3352) and the subset of proteins that did not align to our training set (n=1060). B) The scatterplot contains alignment metrics of antimicrobial resistance proteins from NDARO that were correctly classified by DeepSEA (5654 out of 5959 proteins). The identity cutoff values were retrieved from Arango's work (Arango-Argoty et al., 2018) and the min bitscore value from Pearson's paper about protein homology (Pearson, 2013) C) DeepSEA classification of the remaining 233 proteins that did not align to our training set. D) Count of beta-lactam misclassified as NonR by DeepSEA.
